# Novel frameshift variant in *MYL2* reveals molecular differences between dominant and recessive forms of hypertrophic cardiomyopathy

**DOI:** 10.1101/790196

**Authors:** Sathiya N. Manivannan, Sihem Darouich, Aida Masmoudi, David Gordon, Gloria Zender, Zhe Han, Sara Fitzgerald-Butt, Peter White, Kim L. McBride, Maher Kharrat, Vidu Garg

## Abstract

Hypertrophic cardiomyopathy (HCM) is characterized by enlargement of the ventricular muscle without dilation and is often associated with dominant pathogenic variants in cardiac sarcomeric protein genes. Here, we report a family with two infants diagnosed with infantile-onset HCM and mitral valve dysplasia that led to death before one year of age. Using exome sequencing, we discovered that one of the affected children had a homozygous frameshift variant in *Myosin light chain 2* (MYL2:NM_000432.3:c.431_432delCT: p.Pro144Argfs*57;MYL2-fs), which alters the last 20 amino acids of the protein and is predicted to impact the C-terminal EF-hand (CEF) domain. The parents are unaffected heterozygous carriers of the variant and the variant is absent in control cohorts from gnomAD. The absence of the phenotype in carriers and infantile presentation of severe HCM is in contrast to HCM associated with dominant *MYL2* variants. Immunohistochemical analysis of the ventricular muscle of the deceased patient with the MYL2-fs variant showed marked reduction of MYL2 expression compared to an unaffected control. *In vitro* overexpression studies further indicate that the MYL2-fs variant is actively degraded. In contrast, an HCM-associated missense variant (MYL2:p.Gly162Arg) and three other MYL2 stopgain variants that lead to loss of the CEF domain are stably expressed. However, stopgain variants show impaired localization suggesting a functional role for the CEF domain. The degradation of the MYL2-fs can be rescued by inhibiting the cell’s proteasome function supporting a post-translational effect of the variant. *In vivo* rescue experiments with a *Drosophila MYL2*-homolog (*Mlc2*) knockdown model indicate that neither MYL2-fs nor MYL2:p.Gly162Arg supports regular cardiac function. The tools that we have generated provide a rapid screening platform for functional assessment of variants of unknown significance in *MYL2*. Our study supports an autosomal recessive model of inheritance for *MYL2* loss-of-function variants and highlights the variant-specific molecular differences found in *MYL2*-associated cardiomyopathies.

**Author Summary:** We report a novel frameshift variant in *MYL2* that is associated with a severe form of infantile-onset hypertrophic cardiomyopathy. The impact of the variant is only observed in the recessive form of the disease in the proband and not in the parents who are carriers of the variant. This is in contrast to other dominant variants in *MYL2* that are associated with cardiomyopathies. We compared the stability of this variant to that of other cardiomyopathy associated *MYL2* variants and found molecular differences in the disease pathology. We also show different protein domain requirement for stability and localization of MYL2 in cardiomyocytes. Further, we used a fly model to demonstrate functional deficits due to the variant in the developing heart. Overall, our study shows a molecular mechanism by which loss-of-function variants in *MYL2* are recessive while missense variants are dominant. We highlight the use of exome sequencing and functional testing to assist in the diagnosis of rare forms of diseases where pathogenicity of the variant is not obvious. The new tools we developed for in vitro functional study and the fly fluorescent reporter analysis will permit rapid analysis of *MYL2* variants of unknown significance.

## Introduction

Hypertrophic cardiomyopathy (HCM) is characterized by thickening of the ventricular walls in the absence of a cardiac or metabolic disorder that could account for the hypertrophy [1–4]. It affects 1 in 200-500 individuals and has a strong genetic component [2, 3, 5–7]. It is a major cause of premature sudden cardiac death (SCD), with significant variability in the penetrance and onset of the disease [2, 5, 8]. The majority of HCM patients are asymptomatic, while some display exercise intolerance and progressive heart failure. The ventricular chamber in HCM patients is reduced or normal in size but displays a characteristic inability to properly relax during diastole leading to progressive loss of cardiac function [3, 4, 8–10]. The progressive disease is marked by a disorganized myocyte array in the heart, with significant fibrosis of ventricular walls [7, 11–13]. Consistent with the myofibrillar disarray, about 60% of the cases of HCM have a genetic variant that is associated with the genes encoding the cardiac sarcomeric complex [4, 7, 14–16]. Of these sarcomeric genes, *MYH7, MYBPC3, TNNI3,* and *TNNT2* account for a majority of the variants [13, 16–18]. Identification of novel variants in these genes and other HCM-associated genes has increased dramatically due to the advancement of high throughput genome and exome sequencing technologies [19–23]. However, the advancement of variant identification has also increased the number of potential sequence variants that could contribute to the disease in each individual, confounding the ability to assign pathogenicity to an individual variant [24–26]. While computational methods can predict the damaging effect of a variant and assist in prioritizing variants [27–32], the current consensus on the determination of pathogenicity is dependent on the identification of multiple patients showing similar symptoms and genetic variants with strong evidence of functional impact. Functional testing is therefore critical for disorders that are associated with rare variants in genes affecting a small subset of the patients [25, 33, 34]

Variants in the gene *MYL2* are associated with <5% of cases of HCM [5, 35]. *MYL2* encodes the *Myosin regulatory light chain* expressed in the ventricular heart and slow-twitch skeletal muscles [36]. Even though *MYL2* is considered as a candidate gene for HCM and a handful of pathogenic variants have been described in familial cases and population screens [23, 37–42], the lack of functional testing has prevented the designation of pathogenicity in many cases [43]. Due to the variable penetrance and onset of the HCM phenotype, the pathogenicity of heterozygous *MYL2* loss-of-function variants is hard to assess and have not been directly compared to pathogenic missense variants.

Human *MYL2*, together with the essential light chain (encoded by *MYL3*), stabilizes the ‘lever arm’ of the Myosin head [44]. Human MYL2 is a 18.8 kDa protein with three major domains: A single Ca^2+^-binding EF-Hand domain at the N-terminus, and two EF-Hand like domains in the C-terminus. The N-terminal region also carries a Serine residue (Ser15) which is phosphorylated by Myosin light chain kinase (MLCK) in response to Ca^2+^-mediated activation. This phosphorylation, in turn, modulates the Ca^2+^-Tropomyosin-Troponin dependent activation of Myosin motors in skeletal and cardiac muscles [36]. Phosphorylation of MYL2 by MLCK shifts the equilibrium of the cyclic Myosin head binding to thin filament towards a bound state. This may be due to the increase in the number of cross-bridges between the myosin head and the actin filament and/or through the shift in the average position of the myosin head away from the thick filament and towards the thin filament [36, 45–49]. Correspondingly, variants in the N-terminal EF-hand domain of MYL2 that have defects in Ca^2+^ binding or MLCK phosphorylation are associated with HCM [36, 47]. Several other missense variants that have been identified in both familial and sporadic cases of HCM have been tested in vitro and animal models [48, 50–55].

In mice and zebrafish, loss of the ventricular isoform of the regulatory light chain (*Mlc-2v* in mice and *Myl7* in zebrafish) leads to embryonic lethality with defects in ventricular sarcomere assembly. This indicates the necessity of the regulatory light chain in heart development [56, 57]. On the other hand, mice harboring Mlc-2v phosphorylation mutations or loss of *MLCK* develop to adulthood but die due to heart failure due to dilated cardiomyopathy (DCM) and cardiac hypertrophy with fibrosis [58]. This suggests that in mouse, *Mlc-2v* point mutations are not complete loss-of-function alleles but are hypomorphic alleles. In keeping with this observation, phosphorylation mutants in *Drosophila* regulatory light chain (*Mlc2*) display defect in skeletal muscle organization and flight performance defects in adults while nulls show recessive embryonic lethality [59]. It is worth noting that in mice, heterozygous loss of ventricular regulatory light chain (RLC) does not change the protein level or cardiac function, suggesting that compensatory mechanisms may exist that maintain the level of RLC in these mice [36]. Mice harboring an HCM-associated missense variant MYL2:p.E22K failed to display sarcomeric disarray or changes in cardiac function as measured by echocardiogram [60], while a different mouse model harboring a DCM variant MYL2:p.D94A displays DCM-like phenotype similar to that observed in patients [53].

Such discrepancy in mouse models carrying human variants suggests that disease pathogenesis may differ based on the specific *MYL2* variant. Report of recessive mutations observed in a Dutch family, as well as an Italian patient with Type-I hypotrophy in skeletal muscle along with cardiomyopathy further adds to the variability in disease manifestation [61]. In these cases, *MYL2* variants are thought to be loss-of-function variants, affecting individuals as homozygous recessive, or compound heterozygous variants [61]. Heterozygous carriers of the variant were reported to be asymptomatic mirroring observations in mouse models, where the heterozygous loss of Mlc-2v does not cause a functional defect [61].

In this study, we report a novel recessive variant in *MYL2* identified through exome sequencing in a family where multiple infants have died within one year of age and were diagnosed to have HCM and mitral valve dysplasia. The parents, who are heterozygous carriers of the variant, were found to be normal based on the echocardiographic evaluation. Using *in vitro* analysis we demonstrate that the variant is not stable, and using *in vivo* functional analysis we show that these variants are loss-of-function alleles. By comparing this variant to other HCM-associated variants we propose a role for the C-terminal EF-hand domain in determining the stability of the protein. This will inform in the evaluation of new variants in *MYL2* and provide tools to facilitate rapid screening of these variants.

## Results

### Family with multiple infantile-onset hypertrophic cardiomyopathy and premature death

We identified a family with consanguinity in which four children had died before one year of age (Fig 1A). All four children were born at full-term through a Caesarian section and showed no gross developmental defects (Fig 1B). All four showed a rapid decline in health and were treated for various periods in neonatal intensive care units. Two of them, including the proband, showed a marked increase in heart size while all four children displayed hepatomegaly and general hypotonia (Fig 1B, Fig S1A). Echocardiographic evaluation of the proband showed severe biatrial dilatation and biventricular hypertrophy (Fig S1A). The patient also displayed severe mitral valve regurgitation with abnormal thickening of the mitral valve leaflets and pulmonary arterial hypertension. The proband and the siblings died within the first year of life due to refractory cardiogenic shock and cardiorespiratory arrest. Post-mortem examination of the proband’s heart confirmed severe biatrial dilatation, significant biventricular hypertrophy with small ventricular cavities and severe mitral valve dysplasia (Fig 1C). The father (37 years old) and the mother (27 years old) of the proband, who are consanguineously related (Fig 1A), were asymptomatic from a cardiac standpoint as evaluated by an echocardiogram. While the similarities in the symptoms between the siblings indicated an underlying genetic cause for the disorder, the absence of a family history of cardiac problems confounded a clear diagnosis. Due to the remarkably early onset of HCM in the proband, as well as mitral valve abnormalities, we decided to perform genomic analysis to identify potential disease-contributing variants.

**Fig 1.**
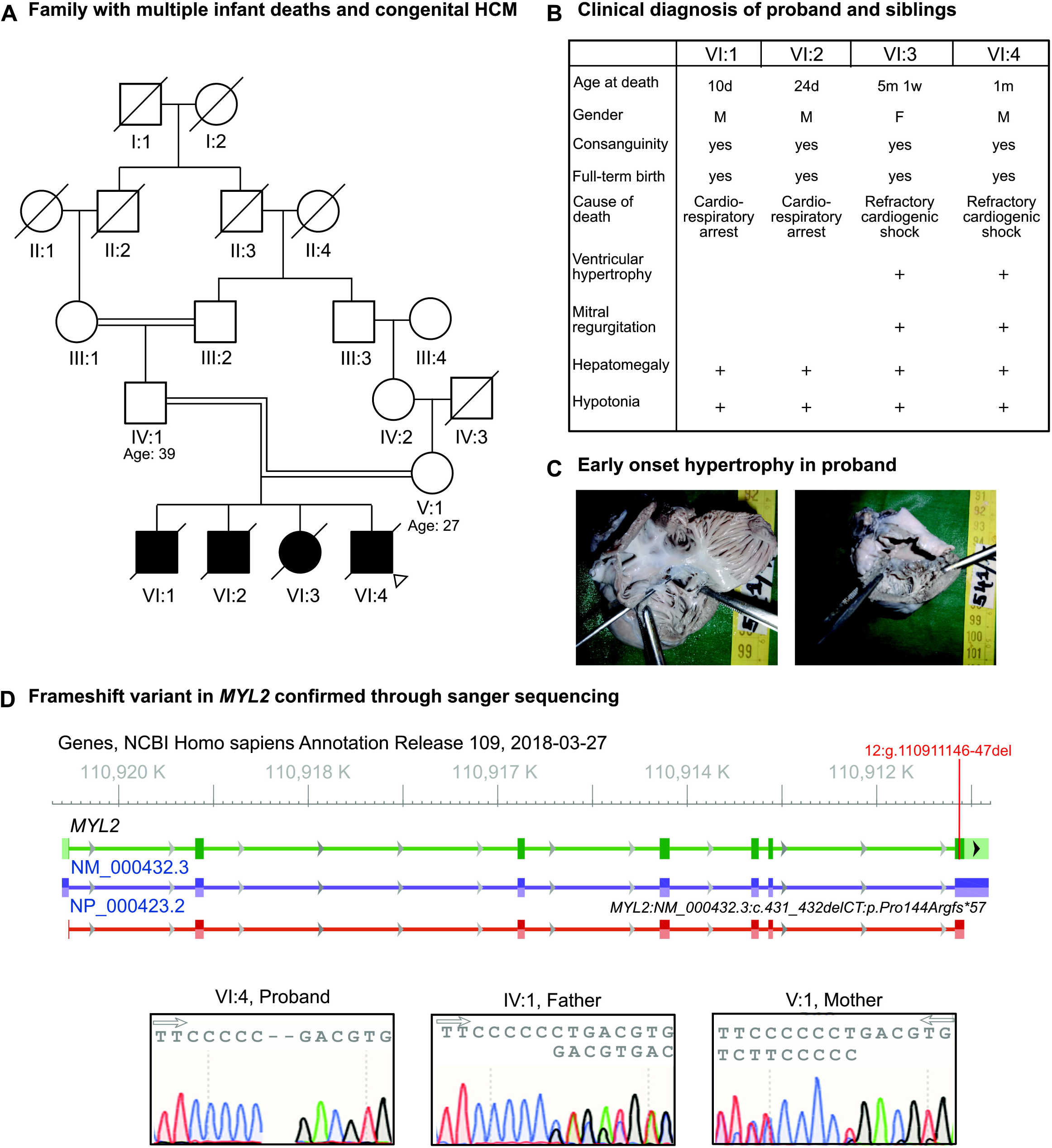
Identification of novel frameshift variant in a proband with infantile hypertrophic cardiomyopathy. (A) Pedigree of the family with multiple infant deaths due to early-onset cardiac disease which also shows consanguinity and age of the parents. (B) Table summarizing key clinical findings of the four siblings. (C) Post-mortem heart dissection of the proband showing severe biatrial dilatation along with hypertrophy of the right ventricle (left) and severe mitral valve dysplasia and hypertrophy of the left ventricle (right) with small ventricular cavities. (D-top) The genomic locus of the *MYL2* gene with the frameshift variant identified in the proband (highlighted in red) in the last exon of the gene. (D-bottom) Sequence chromatograms from Sanger sequencing of proband showing homozygosity of the frameshift variant. The chromatograms of the parents show heterozygosity of the frameshift variant.

### Whole exome sequencing to identify variants and prioritization of variants for functional analysis

We performed exome sequencing on the proband, the mother and the father. Sequencing data were analyzed using our previously published pipeline, Churchill, for calling variants [62]. Variants were prioritized using minor allele frequency (<0.001) and damaging effect prediction by five out of the seven algorithms to filter variants (Fig S1B) [27–32]. This approach resulted in the identification of one *de novo* heterozygous variant in *Olfactory Receptor Family 7 Subfamily C Member 1* (*OR7C1*:NM_198944.1:c.335delA:p.Asn112fs) and a homozygous variant in *Myosin light chain 2* (*MYL2*:NM_000432.3:c.431_432delCT:p.Pro144Argfs*57) (Fig 1D). *OR7C1* encodes an olfactory receptor protein that is not expressed in the heart. Therefore, we focused on the *MYL2* variant (*MYL2-fs*). We confirmed the presence of the homozygous variant in the proband and that the parents are heterozygous carriers using Sanger DNA sequencing of the genomic region (Fig 1D). The dinucleotide deletion in the last exon of *MYL2* is predicted to cause a frameshift variant of the protein that affects the last 20 amino acids in the C-terminal EF-Hand domain. Moreover, the variant extends the reading frame of *MYL2* into the 3′ UTR, leading to the addition of 36 amino acids to the C-terminal end. This *MYL2-fs* variant was not found in control populations in gnomAD, supporting the possibility of its association with this rare disorder.

### Myocyte disarray, fibrosis, and reduction in the expression of MYL2 in the proband’s ventricular muscle

To test the consequence of the variant, we examined the histopathology of the myocardium in the proband’s ventricular muscle. Consistent with HCM diagnosis, the patient’s ventricular muscle displayed myocyte disarray and a substantial increase in fibrotic tissue as evidenced by the H&E and Masson’s trichrome staining respectively (Fig 2A). Next, we examined the expression level of MYL2 using immunohistochemistry. We observed a significant reduction in the MYL2 protein levels in the proband’s ventricular muscle compared to an unaffected control sample. In contrast, the expression level of a different sarcomeric protein, cardiac troponin I (TNNI3), in the proband was similar to control (Fig 2B). Also, *MYL2* mRNA could be detected in the proband’s ventricle using RT-PCR (Fig 2C). This suggests that the frameshift variant adversely affects the levels of MYL2 protein in the proband.

**Fig 2.**
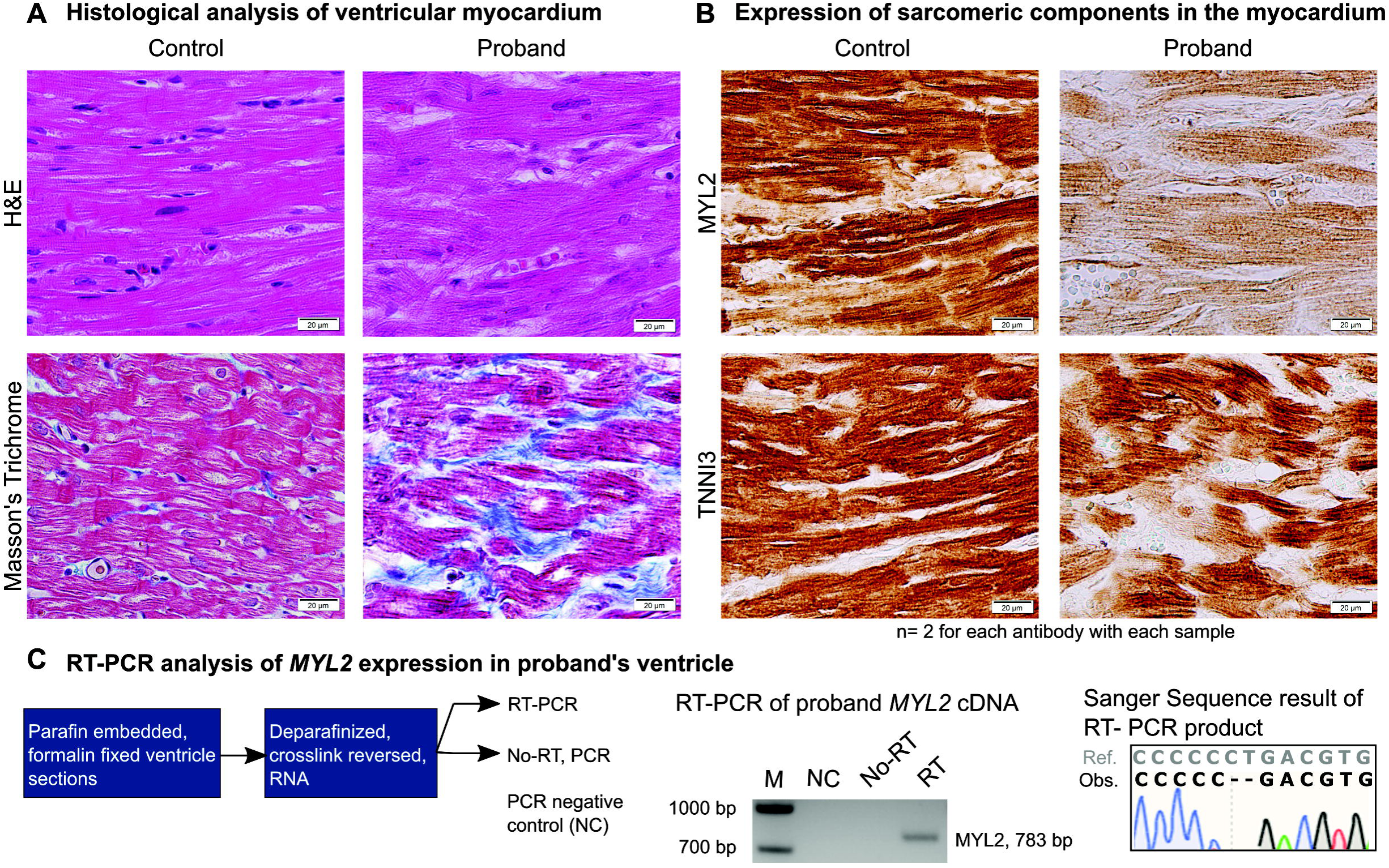
Analysis of proband’s ventricular myocardium shows characteristic features of hypertrophic cardiomyopathy (HCM) and molecular consequence of the variant. (A) Histochemical analysis of ventricular myocardium of control and proband’s samples using H&E staining showing myocyte disarray in the proband. Masson’s trichrome staining of the proband’s ventricular myocardium showing increased blue-stained fibrosis. (B) Immunohistochemical analysis of MYL2 expression in control and proband’s ventricular myocardium showing a remarkable reduction in the expression of MYL2 in the proband. Cardiac troponin I (TNNI3) is equally expressed in the control and the proband myocardium; the non-stained areas correspond to fibrosis. (C) Flow diagram showing the steps used to analyze *MYL2* mRNA expression in the proband ventricular myocardium. PCR product corresponding to the expected size of *MYL2* mRNA is detected in the proband, and not in the negative controls (NC, No-RT). Sanger sequencing of this product shows the expression of *MYL2* mRNA with the dinucleotide deletion.

### *In vitro* functional testing of the stability of MYL2 variants

We hypothesized that the reduction in the levels of MYL2 in the proband’s ventricular muscle is due to the instability of the protein product with frameshift mutation. To test this hypothesis, we compared the stability of the wildtype and frameshift variants overexpressed in rat cardiomyoblast cells (H9c2 cells) [63]. We generated an EGFP-tagged human *MYL2* cDNA construct that also expresses mCherry, permitting simultaneous evaluation of mRNA stability and protein stability. Using this construct, we tested the stability of MYL2-frameshift variant along with three other loss-of-function, stopgain alleles and a missense allele mapping to the C-terminal EF-hand domain (MYL2:p.G162R) reported in ClinVar (Fig 3A). While it is likely that the stopgain mutations (not found in the last exon of MYL2) will be degraded through non-sense mediated decay (NMD) when expressed from the genomic loci, the cDNA overexpression analysis allowed us to examine the effect of loss of different domains of the MYL2 protein. Using immunoblot analysis, we observed that overexpression of MYL2-fs variant was significantly reduced compared to truncation variants and the missense variant (Fig 3B, Fig S3B). However, there was no significant difference in the mCherry signal between these constructs (Fig 3B). We noticed that truncated variants did not localize in a pattern similar to the wild type MYL2 protein, which showed strong localization along the cytoskeleton of the cells (Fig 3C). The fs variant, however, was not detected in H9c2 cells compared to the wildtype control. Once again, there was no significant difference in the production of mCherry from fs variant constructs, suggesting that the read-through into the 3′ UTR does not destabilize the mRNA (Fig 3C). This prompted us to examine the mode of degradation of the MYL2-fs variant. We examined if the MYL2-fs variant is degraded by the proteasome machinery by treating the cells with MG-132, a commonly used pharmacological inhibitor of the proteasome. Indeed, when the H9c2 cells were treated with MG-132 after transfection with the MYL2-frameshift construct, the GFP signal corresponding to MYL2-fs was recovered (Fig 3D). However, it is worth noting that the MYL2-frameshift variant appears to be aggregating and has a weak affinity towards the cytoskeleton (Fig 3D). The binding of the MYL2-fs to myosin head may be affected as the modified residues are found close to the lever arm (Fig S3A). The aggregation suggests that the instability of the MYL2-fs variant might be due to misfolding or changes in the biochemical properties of the protein. Together, the *in vitro* experiments indicate that the MYL2-fs variant is not stable, and the loss of stability is due to proteasome-mediated degradation of the translated product and not due to changes in mRNA stability.

**Fig 3.**
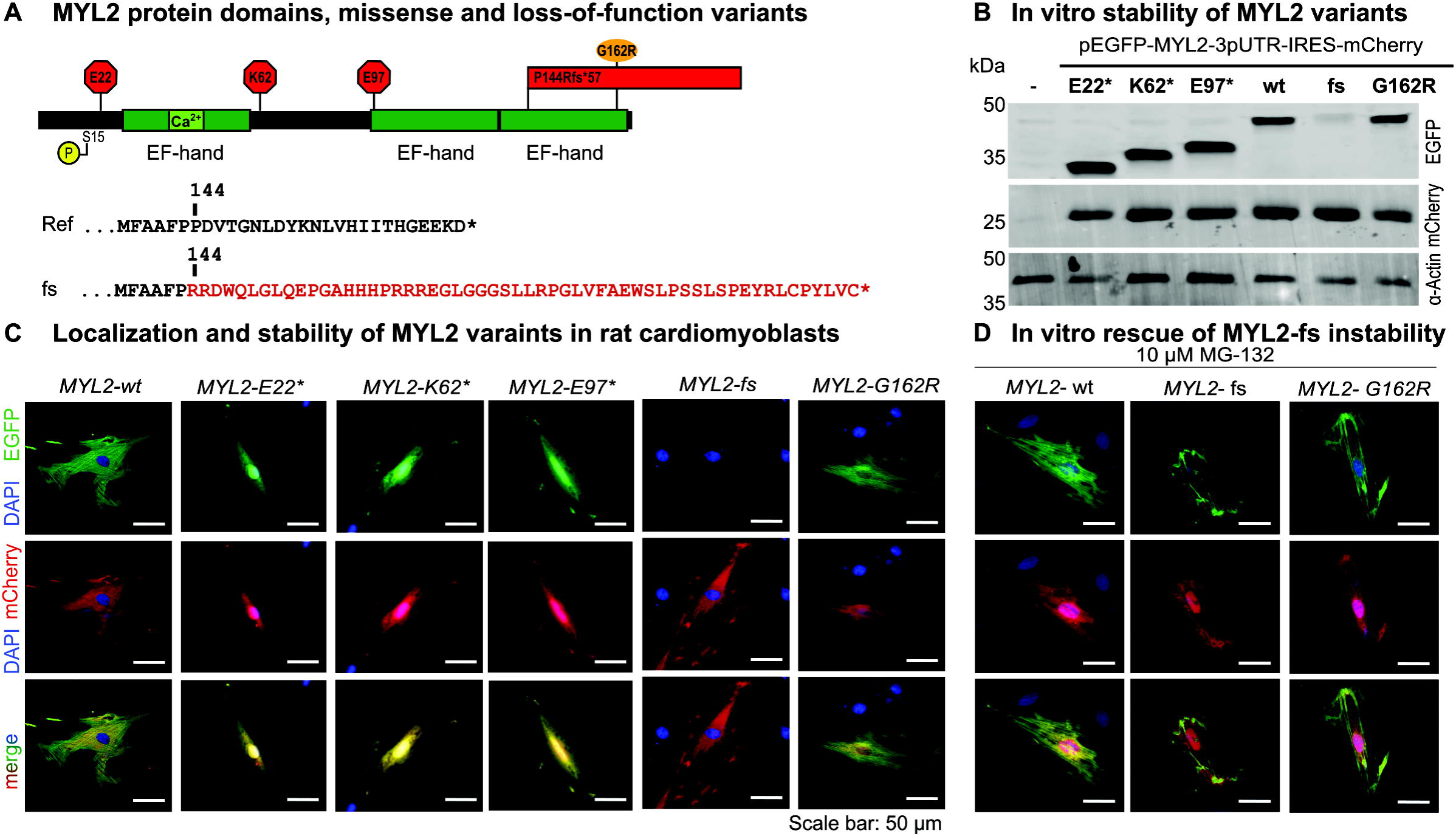
*In vitro* analysis of stability and localization of MYL2 variants. (A) Primary structure of MYL2 showing N-terminal & C-terminal EF-hand domains, Serine 15 residue that is target of MLCK phosphorylation, three stopgain variants (red octagons), the frameshift variant identified in the proband (red block showing extension of C-terminal end) and missense variant associated with HCM (orange oval) that maps the C-terminal domain. The frameshift mutation changes the canonical MYL2 sequence (‘Ref.’, black text) from residue 144 onwards, modifying the last 20 amino acids of the protein in addition to adding 36 non-canonical amino acids to the C-terminus (‘fs’, red text). (B) Western blot image showing overexpression of EGFP-tagged MYL2 wt and variant cDNAs. The stability of the frameshift variant is significantly affected while other loss-of-function variants did not show a significant reduction in stability. mCherry signal analyzed using western blot does not show any significant changes between MYL2 wt and variant sequences. Loading controls: mCherry and Actin. (C) Immunofluorescence analysis of EGFP tagged MYL2 wt and variants (green) co-overexpressing mCherry from the same transcript (red) in H9c2 cells. EGFP tagged MYL2-wt localizes to the cytoskeleton, while stopgain variants (E22*, K62*, E97*) do not localize to the cytoskeleton and are observed in a diffuse pattern. EGFP signal from the frameshift variant (fs) is significantly reduced. Localization of the missense (G162R) variant is in a pattern similar to the wild type. mCherry signal from the transfected cells does not change between variants. (D) Immunofluorescence images of EGFP tagged MYL2 wt and variants transfected and MG-132 treated H9c2 cells showing rescue of MYL2-fs variant signal following MG-132 treatment. Scale bar indicates 50 uM length. Quantitation of signal from the western and immunofluorescence are provided in Fig S3.

### *In vivo* functional analysis using a *Drosophila* model of Myosin light chain knockdown in the dorsal tube

To test if any residual expression of the MYL2-fs variant could support myosin light chain function, we examined the ability of human wildtype *MYL2* and variants to rescue the loss of *Drosophila* myosin light chain (*Mlc2;* Fig S4) expressed in the heart. We used *Drosophila Hand-GAL4* driver to knock down the expression of *Drosophila Mlc2* using transgenic RNAi lines (Fig 4A). This led to a significant loss in developmental viability, as measured by percent of observed adult survivors compared to the expected Mendelian frequency of the knockdown genotype (Fig 4C). In the knockdown background, using the same *Hand-Gal4* driver, we overexpressed the human *MYL2* cDNA or the variant carrying cDNAs to test the ability of the human *MYL2* to functionally substitute fly *Mlc2* (Fig 4A). We observed that the developmental lethality caused by cardiac knockdown of *Mlc2* was partially rescued by the wildtype human *MYL2* overexpression, while the overexpression of the frameshift variant identified in our proband or the MYL2-G162R missense variant failed to rescue the phenotype. To examine the specific effect of the knockdown and the variant overexpression in the heart, we focused on the cardiac function in the third instar larvae during the fly development. We used a membranous mCherry reporter (CD8::mCherry) to track the rhythmic contraction of the *Drosophila* heart between the posterior denticle belts (A7-A8) (Fig 4B, Movie S1). Using this reporter driven by the *Hand-GAL4,* we measured fractional shortening of the heart and compared it between different genotypes (Fig 4D, Movie S1-S5). We observed that loss of *Mlc2* due to RNAi-mediated knockdown lead to a decrease in fractional shortening, which was again partially rescued by the wildtype human *MYL2* cDNA overexpression. As seen with the developmental lethality, there was no rescue observed with the MYL2-fs or the MYL2-G162R variant in terms of fractional shortening (Fig 4 D, Movie S1-S5). Partial rescue in both experiments can be attributed to sequence differences between the human and *Drosophil*a homologs. *Drosophila* Mlc2 N-terminal region has additional sequences that have been shown to be needed for *Mlc2* function [64]. Nevertheless, the *in vivo* analyses suggest that the frameshift variant and missense variant are functionally different from the wildtype human *MYL2*, and therefore will not support adequate cardiac function.

**Fig 4.**
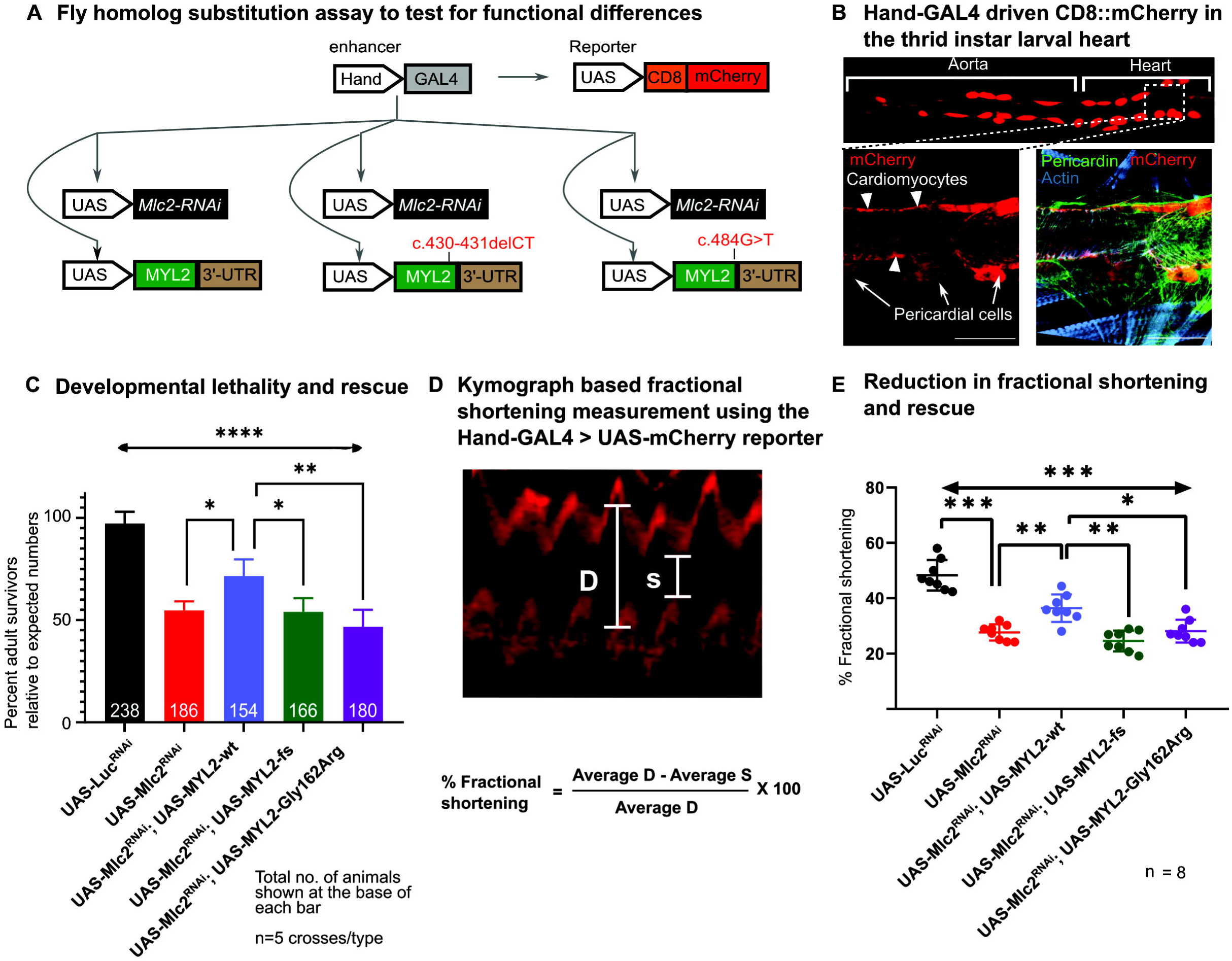
*In vivo* functional analysis of MYL2 variants using *Drosophila Mlc2* substitution assay in the heart. (A) Schematics showing Hand-enhancer driven GAL4 (Hand-GAL4) used to knockdown *Mlc2* expression using RNAi in the heart, and to overexpress human MYL2-wt and variant cDNAs for functional substitution. In parallel, the Hand-GAL4 drives the expression of UAS-CD8::mCherry which facilitates visualization of the heart. (B-top) Immunofluorescence image of the *Drosophila* heart from UAS-CD8::mCherry; Hand-GAL4 animals showing expression of mCherry in the heart chambers and aorta. (B-bottom) Magnified image of the A7-A8 posterior denticle band region showing expression of mCherry in both cardiomyocytes as well as pericardial cells. Merged imaged on the right shows cardiac cells marked by Pericardin expression (green), mCherry driven by Hand-GAL4 (red) and Phalloidin staining of actin (blue) marking both cardiomyocytes and skeletal muscle. No expression of mCherry was detected in the skeletal muscle. (C) Histogram showing developmental lethality due to *Mlc2* RNAi knockdown using Hand-GAL4 and rescue by overexpression of human MYL2-wt and variants. Partial rescue of the lethality is observed with wildtype *MYL2* cDNA overexpression while no significant difference is observed for the tested variants. Percentage of adult survivors calculated based on the fraction of adults with the obtained desired genotype compared to Mendelian expected ratio for each cross. Significance analyzed using Brown-Forsythe and Welch ANOVA across all samples followed by multiple comparisons using Games-Howell multiple comparison test. Significance indicated using * adj. P-Value < 0.05, ** adj. P-Value < 0.01, **** adj. P-Value < 0.0001. (D) Kymograph image showing a rhythmic pattern of cardiac contraction observed using UAS-CD8::mCherry reporter driven by Hand-GAL4. Diastolic width is shown by ‘D’, and systolic width by ‘S’. The formula used to calculate fractional shortening is shown below the kymograph. (E) Interleaved scatter plot showing fractional shortening across different genotypes (Movies S1-S5). Fractional shortening was significantly reduced due to *Mlc2* RNAi knockdown which is partially rescued by overexpression of wild-type human *MYL2*. The missense and fs variants showed no significant improvement in fractional shortening compared to the wild type. Significance was calculated using two-way ANOVA followed by Tukey’s multiple comparison tests. Multiplicity adjusted P-Values of Tukey’s comparison are indicated as: * adj. PValue < 0.05, ** adj. PValue < 0.01, *** adj. PValue < 0.001, **** adj.PValue < 0.0001.

## Discussion

Using next-generation exome sequencing methodology and our variant prioritization pipeline, we have identified in the gene *MYL2*, a rare and novel frameshift variant in a family with multiple infant deaths and early onset HCM. Consistent with this diagnosis, we found that the patient’s ventricular myocardium shows a marked increase in fibrosis and myocyte disarray. The frameshift variant leads to a significant reduction in MYL2 in the ventricular myocardium. This variant does not destabilize the mRNA, as we could detect the transcript in the patient’s ventricle as well as through the observation of mCherry reporter in the in vitro stability assay. The *in vivo* functional testing further demonstrated that any residual MYL2-fs variant present will not be able to adequately support cardiac function. The *in vitro* and *in vivo* tools generated in this study will permit rapid analysis of functional differences in variants of unknown significance in *MYL2*.

These functional testing results shed light on the molecular pathology of the MYL2-fs variant and recent reports of other recessive MYL2 variants leading to early-onset cardiomyopathy [61, 65]. The instability of the MYL2-fs variant suggests that any negative effects of MYL2-fs variant on the myosin function is not likely to be dominant. This conclusion is also supported by several loss-of-function variants reported in gnomAD, rendering the probable likelihood of intolerance for *MYL2* to be zero [66]. This brings to the fore two aspects of MYL2 loss-of-function variants: 1) Loss-of-function variants may not contribute to HCM due to haploinsufficiency. 2) There is a strong need to test individual variants to understand their mode of pathogenicity in order to develop a personalized therapeutics.

The development of the ventricular myocardium in the absence of *MYL2* might be supported by the increase in the atrial light chain *MYL7* expressed in the embryonic ventricle, as observed in the mouse models [56]. Whether such a response exists in dominant *MYL2*-variant associated myopathies remains to be investigated. This is important due to the observation that missense variants display reduced capacity to support myosin function *in vitro* [67]. However, the missense variants can still impart a negative effect on myosin function by affecting myosin contractility [48]. This may not be in the case in the stop-gain mutants that are likely to be degraded through NMD or frameshift variants that result in destabilized protein. The observation that loss-of-function alleles display recessive inheritance while missense variants display dominant inheritance in HCM is supported by animal model analyses [56, 57] and is similar to genetic analysis of other sarcomeric genes [68]. However, the possibility exists that other variants in the patient could modulate the instability of the MYL2-fs variant, such as reduced proteasome function, that could easily stabilize the protein. Therefore, there is a need to identify modifiers and detect variants in these genes in future patients with MYL2 loss-of-function variants to determine the risk of HCM inheritance. Unlike the missense variants, where allele-specific silencing approach may provide a viable solution [69], the treatment and management of recessive form of HCM will require an *in utero* diagnosis of the disorder and may involve gene therapy using a functional copy of *MYL2*. While this may not be immediately feasible, the diagnosis and genetic counseling of carriers need to be considered.

## Materials and Methods

### Patient recruitment and DNA isolation

Informed consent was obtained from the Parents of the proband as per Institutional Review Board guidelines at University of El Manar, Tunisia. DNA was isolated using standard procedures.

### Whole-exome sequencing

Exome libraries were constructed using the Agilent SureSelectQXT Target Enrichment System for Illumina Multiplexed Sequencing Protocol (Agilent Technologies, CA). DNA libraries were captured with the Agilent Clinical Research Exome Kit. Paired-end 150 base pair reads were generated for exome-enriched libraries sequenced on the Illumina HiSeq 4000 to a targeted depth of 100× coverage.

### Variant identification, prioritization, and confirmation

The primary and secondary variant analysis was performed as described in our previous studies. The analysis of the sequencing data was conducted using the Churchill pipeline [62], in which the data was aligned to GRCh37 using BWA mem, deduplicated using samblaster, and variants were jointly called across all samples using GATK’s HaplotypeCaller. SnpEff, a software tool to annotate genetic variation, was used along with custom in-house scripts to provide mutation and gene information, protein functional predictions and population allele frequencies. Common variation occurring at >0.1% minor allele frequency in the population was excluded. Variants outside of coding regions (defined as >4 base pairs from an exon splice site) and exonic variants coding for synonymous single nucleotide polymorphisms were also dropped. *In silico* analysis was performed using algorithms to predict the pathogenicity of identified sequence variants. The following prediction software was used to analyze the rare variants in candidate CHD genes identified through WES: SIFT, GERP++, Polyphen2 Complex, Polyphen2 Mendelian, MetaSVM, MetaLR, and CADD [27–32]. Variants were further filtered based on the expression of the impacted human gene in the developing heart using publicly available single-cell data [70]. *MYL2* genomic region flanking the identified variant was amplified using MYL2.gen.For and MYL2.gen.Rev primers and the presence of the variant was confirmed using Sanger sequencing method.

### Total RNA isolation from patient ventricle and RT-PCR

Four 7 micron sections of the left ventricle were used to isolate total RNA using the Quick-RNA^TM^ FFPE kit (Zymo Research). 1 µg of total RNA was used to generate cDNAs using the SuperScript™ VILO™ cDNA Synthesis Kit. MYL2 cDNA was amplified using the primers FP.MYL2.BglII and RP.MYL2-3p-UTR.XhoI and the PCR product was sequenced to confirm the expression of MYL2-fs variant.

### Plasmids constructions

Human *MYL2* cDNA (Refseq ID NM_000432.3) in the pDNR-LIB plasmid was obtained from Harvard Plasmid Database. *MYL2* coding region with the 3′ UTR sequence was amplified using PCR with primers FP.MYL2.BglII and RP.MYL2.EcoRI, and cloned into *Bgl*II/*Eco*RI site in the pIRES-mCherry plasmid (a gift from Ellen Rothenberg (Addgene plasmid # 80139; http://n2t.net/addgene:80139; RRID: Addgene_80139). Then, *MYL2*-coding region, 3′ UTR sequence and IRES mCherry were excised using BglII/ClaI and introduced in frame with the EGFP tag in a modified pEGFP-C1 plasmid (Clonetech) to generate the pEGFP-MYL2-IRES-mCherry plasmid. This construct, when transfected in cells, leads to the production of an EGFP tagged MYL2 protein (or MYL2 variant protein) and independently translated mCherry from the same transcript. The EGFP tag distinguishes the overexpressed cDNA from rat *Myl2,* while the independently produced mCherry acts as a readout of mRNA stability and also serves as a transfection control allowing direct comparison of cellular levels of the MYL2 protein variants. For *Drosophila* transgenic experiments, untagged MYL2 coding region with the 3’UTR was amplified using FP.MYL2.BglII and RP.MYL2-3p-UTR.XhoI inserted using BglII/XhoI in pUASt-attB-exp (a modified pUASt-attB vector with additional restriction sites) to generate pUASt-MYL2-attB. To generate the variants, site-directed mutagenesis was used with pEGFP-MYL2-IRES-mCherry or pUASt-MYL2-attB as the template and using the following primers and Agilent Quickchange II kit [71]. Sequences are provided in Table S1.

MYL2-fs (c.431-432delCT): FP.delCT-431-432.MYL2, RP.delCT-431-432.MYL2

MYL2-G162R: FP.MYL2.G162R, RP.MYL2.G162R

MYL2-E22*: RP.MYL2E22Stop, FP.MYL2E22Stop

MYL2-K62*: FP.MYL2K62Stop, RP.MYL2K62Stop

MYL2-E97*: RP.MYL2E97Stop, FP.MYL2E97Stop

### Cell culture

H9c2 cells [63] were cultured in Dulbecco’s Modified Eagle’s Medium (DMEM) with 4.5 g/L Glucose, 4 mM L-Glutamine, 1 mM sodium pyruvate, and 1.5 g/L sodium bicarbonate (ATCC 30-2002), supplemented with 10% fetal bovine serum, 100 I.U./mL penicillin and 100 (μg/mL) streptomycin at 37 °C incubator with 5% CO_2_. Cells were transfected with 2 µgs of the plasmid with Lipofectamine 3000 reagent with OptiMEM media according to manufacturer’s recommendations. Transfection media were removed five hours post-transfection and replaced by normal growth media. Cells were collected for Immunoblot analysis or Immunofluorescence 48 hours after transfection. For proteasome inhibition, 24 hours post-transfection, cells were treated with 10 uM MG-132 in growth media and incubated for another 24 hours before analysis.

### Fly stocks

*Drosophila* lines were maintained in standard fly food with yeast at 25 °C. The following stocks were obtained from the Bloomington *Drosophila* Stock Center: *UAS-CD8.mCherry* (BDSC 27391), *Hand-enhancer-GAL4* (BDSC 48396), ‘Dm integrase with attP landing site VK37’ (BDSC 24872), *Mlc2^RNAi^* (JF01106, BDSC 31544), Luciferase (firefly)^RNAi^ (BDSC 31603), Multiple balancer stock with mCherry marker (BDSC 76237; CyO, P{sqmCh}2; TM3, P{sqmCh}3, Sb).

### *Drosophila* transgenesis and crossing schemes

UAS-*MYL2*-attB constructs (300 ng/uL) were injected into ‘Dm integrase with landing site VK37’ stock to generate overexpression transgenic stocks using previously described methods with small modifications. F0 injection-survivors were crossed to *yw* animals and transgenic progeny from this cross (F1) were identified using red eye color. Stocks were generated using standard *Drosophila* mating schemes to generate UAS-MYL2 (wt or variant)/Cyo P{sqmCh}2; Mlc2^RNAi^/TM3, Sb, P{sqmCh}3 stocks. These were crossed to UAS-CD8::mCherry/Cyo; GMR88D05-Hand GAL4/TM6, Ubx-LacZ animals and raised at 29 °C to maximize the effect of GAL4 mediated transcription. For developmental lethality, emerging adults irrespective of sex were used to determine percent adult survivors and compared to control crosses where *Luciferease* RNAi was overexpressed using *Hand-GAL4*. For cardiac function analysis, larvae of the desired genotype were identified using mCherry signal in the heart and the absence of muscle mCherry signal from the balancer chromosome.

### Fluorescent reporter based fly cardiogram

Crawling-third instar larvae were collected for each genotype and immobilized on a double-sided tape on a glass slide with the dorsal side of the larvae towards the camera. Videos were collected using an Olympus BX51 microscope with a DP71 camera. Each larva was imaged two times for 15 seconds each with 15 seconds gap. Kymographs of the videos were created, and systolic-diastolic widths of the heart tube were measured using ImageJ across 10 different contractions and averaged between the two videos for each animal. Fractional-shortening was calculated as described in previous studies using the formula: % fractional shortening = (Avg. D – Avg. S) / (Avg. D) *100 where ‘D’ is the diastolic width and ‘S’ is the systolic width [72].

### Western blot

Immunoblots were performed using standard protocols prescribed for LI-COR biosciences method of detection using infra-red dye conjugated secondary antibodies. The following antibodies were used: Chicken anti-GFP (1:1000; Abcam: ab13970), Rabbit anti-mCherry (1:1000; Abcam: ab167453) and Rabbit anti-actin (1:1000; Abcam: ab1801).

### Histology, Immunohistochemistry, and Immunofluorescence

Formalin-fixed and paraffin-embedded ventricular myocardium of the deceased proband from the Department of Embryo-Fetopathology of Tunis and control heart from an unaffected donor of comparable age obtained from the Biospecimen repository at Nationwide Children’s Hospital were sectioned and processed using standard methods. Hematoxylin and Eosin Staining [73] and Masson’s trichrome staining [74] were used to study the histology of the samples. For immunohistochemistry following antibodies were used: Rabbit Anti-Myosin light chain 2 antibodies (1:500, Abcam: ab48003), Rabbit anti cardiac troponin I (TNNI3) (1:500, Abcam ab47003). For immunofluorescence, cells were fixed using 4% paraformaldehyde and processed for immunofluorescence using standard protocols. The following antibodies were used: Chicken anti-GFP (1:500; Abcam: ab13970) and Rabbit anti-mCherry (1:500; Abcam: ab167453). *Drosophila* larvae were dissected and stained with Pericardin antibody (1:100, Developmental Studies Hybridoma Bank: EC11) as described previously.

## Acknowledgments

We thank the Tunisian family members for their participation in this study. We also thank Dr. Amanda Simcox for guidance in *Drosophila* husbandry.

## Supporting information

**Fig S1: Clinical diagnosis and identification of rare variant in siblings.** (A) X-ray radiographs showing hepatomegaly in all four siblings and cardiomegaly in the proband (VI:4) and sister (VI:3). Echocardiograph analysis of the proband showing severe bi-atrial dilation (left and right) and small ventricular cavities with significant septal hypertrophy (left) on planes comparable to post-mortem analysis in Fig 1C. (B) Flow diagram showing the number of genes with variants passing through each filter in our variant prioritization pipeline. Variants are classified into *de novo*, Homozygous and Compound Heterozygous variants before prioritization. Three filters are used to reduce possible sequencing artifacts (Quality score cutoff), common variants (Frequency in general population cutoff), and variants predicted to be benign (Damaging effect prediction cutoff). One *de novo* and one homozygous variant were prioritized using this pipeline.

**Fig S2: Model of Myosin interacting-heads motif (IHM) showing impacted residues.** (A) A model of Myosin interacting head motif showing Myosin heavy chain (blue) and two interacting light chains: Essential light chain (red), Regulatory light chain (green-pink). In the regulatory light chain, residues affected by frameshift variant are shown in pink with the starting residue (Pro144) shown as spheres. The proximity of the variant residues to the IHM suggest that it can affect the binding of the regulatory light chain to the myosin head.

**Fig S3: *MYL2* variants, reporters and in vitro assays.** (A) MYL2 primary structure showing domains and missense variants reported in ClinVar (Cyan bubbles). The frameshift variant is shown as a red box. (B) EGFP-tagged MYL2 overexpression vectors showing various overexpression products that are expected from each construct. (C) Quantitation of immunoblot based stability analysis of *MYL2* variants (left) and Immunofluorescence based stability analysis (right). One-way ANOVA was used to test for significance and multiplicity adjusted P-Values from Tukey’s multiple comparisons are shown. ANOVA ** P-Value < 0.05.

**Fig S4: Conservation of amino acid sequence between MYL2 homologs.** (A) MUSCLE alignment of human MYL2 and homologs from different species show high conservation of residues along the length of the protein.

**Movie S1: Third instar larval heart used as control (GMR-Hand-GAL4> UAS-Luc^RNAi^)**

**Movie S2: Third instar larval heart showing the impact of Mlc2 knockdown (GMR-Hand-GAL4> UAS-Mlc2^RNAi^)**

**Movie S3: Third instar larval heart showing partial rescue of Mlc2 knockdown by human MYL2 wt cDNA overexpression (GMR-Hand-GAL4> UAS-Mlc2^RNAi^, UAS-MYL2-wt)**

**Movie S4: Third instar larval heart showing no rescue of Mlc2 knockdown by human MYL2-fs cDNA overexpression (GMR-Hand-GAL4> UAS-Mlc2^RNAi^, UAS-MYL2-fs)**

**Movie S5: Third instar larval heart showing no rescue of Mlc2 knockdown by human MYL2-G162R cDNA overexpression (GMR-Hand-GAL4> UAS-Mlc2^RNAi^, UAS-MYL2-G162R)**

**Table S1:**
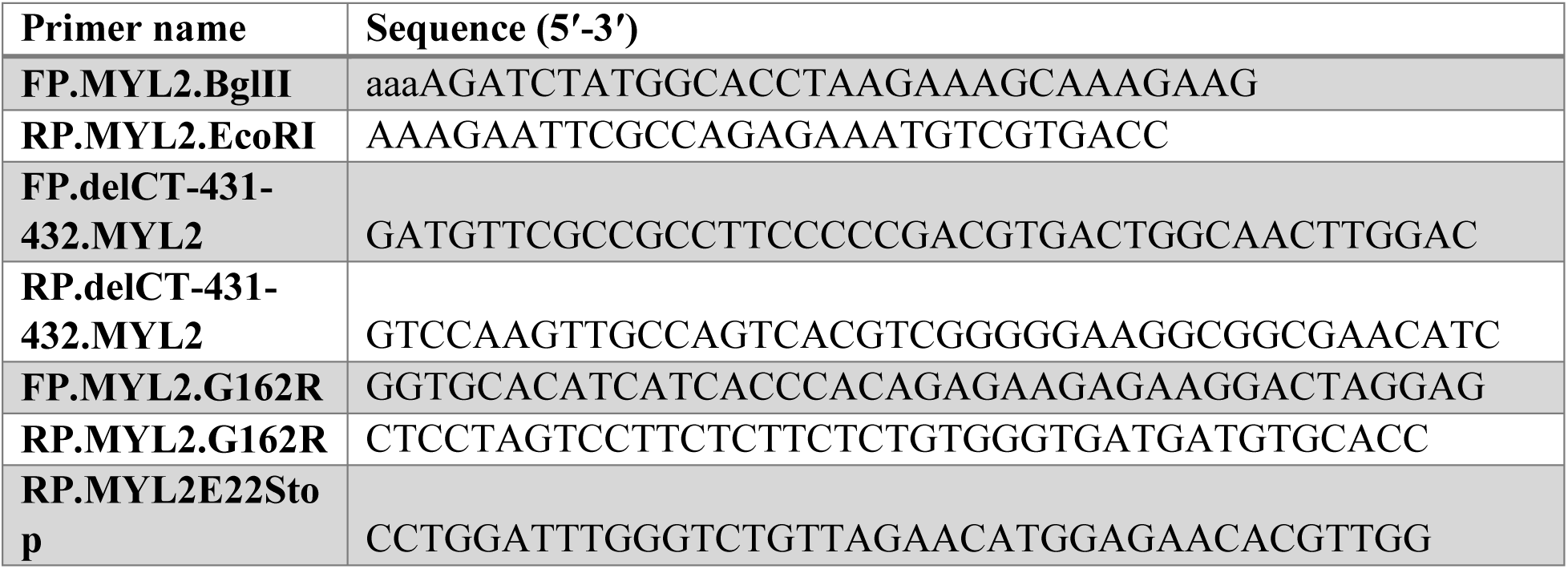
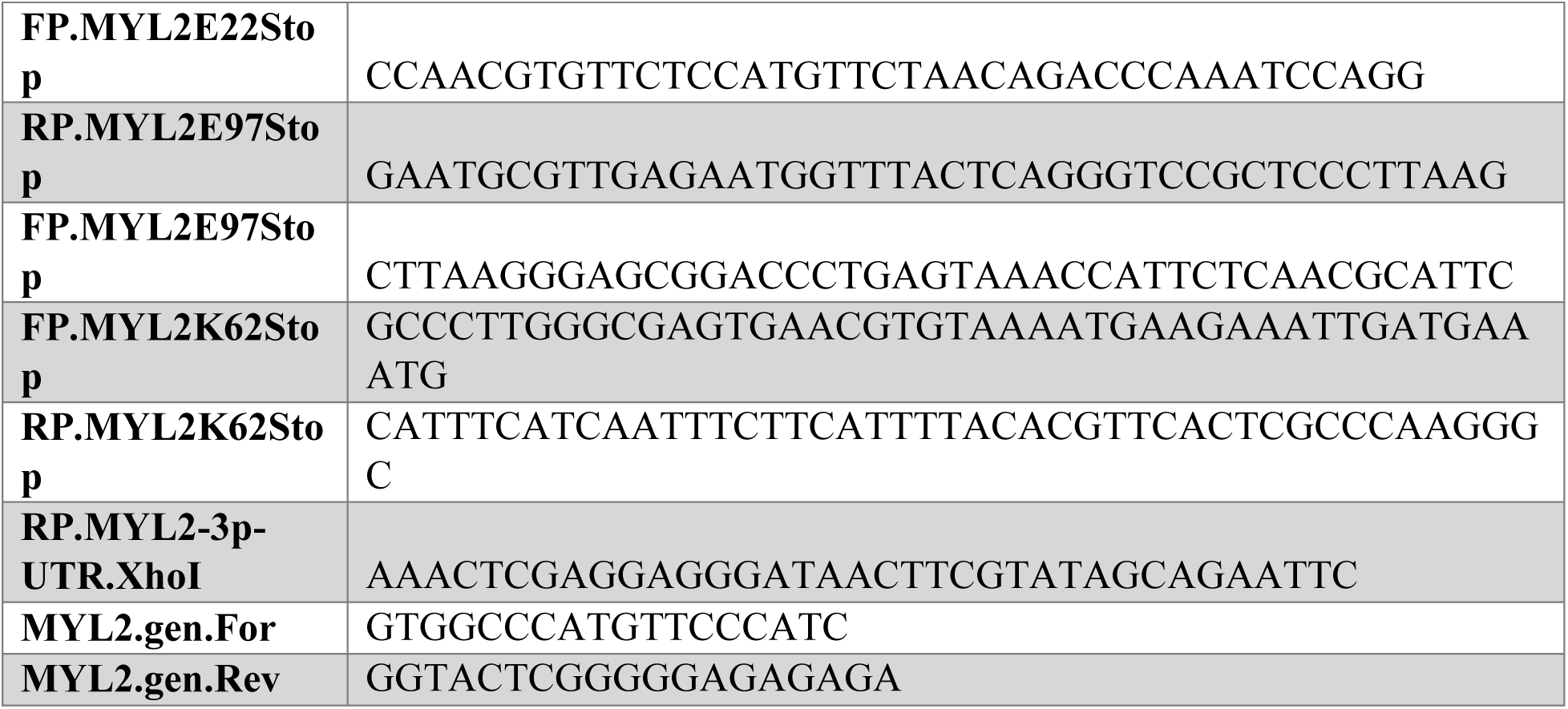
Sequence of primers used in the study

