## Supplementary figures and images for "Novel frameshift variant in *MYL2* reveals molecular differences between dominant and recessive forms of hypertrophic cardiomyopathy"

### Fig. S1

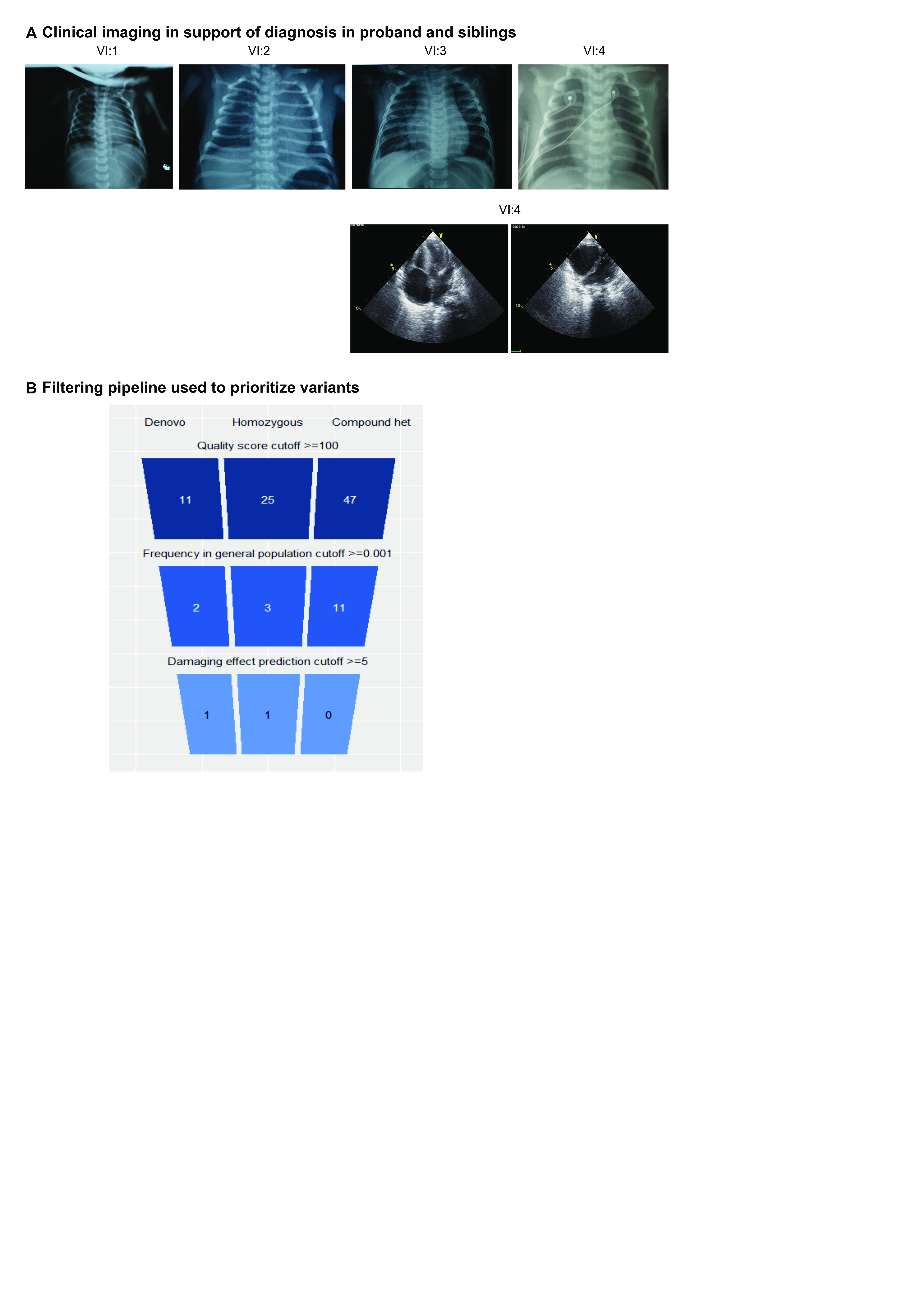

### Fig. S2

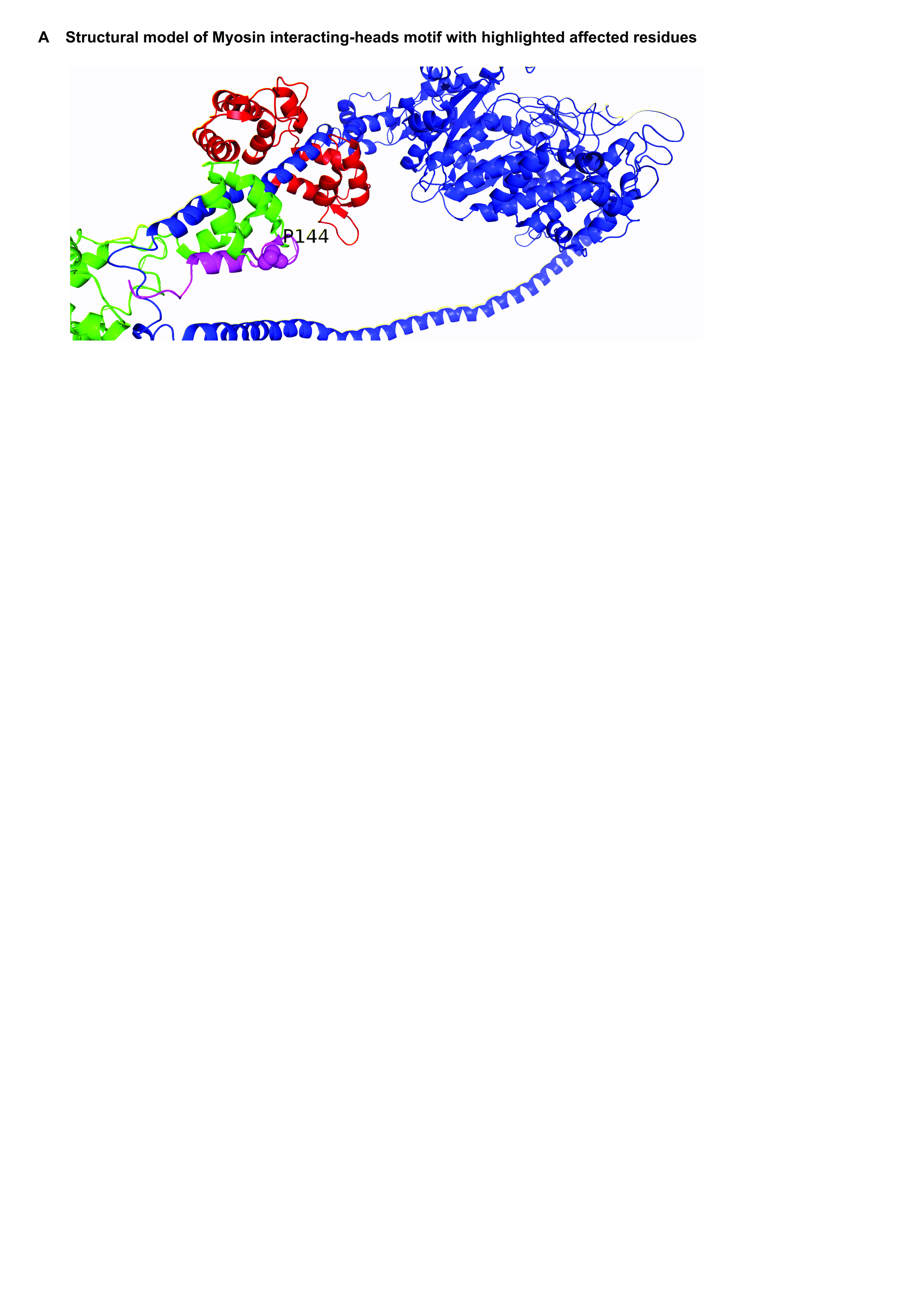

### Fig. S3

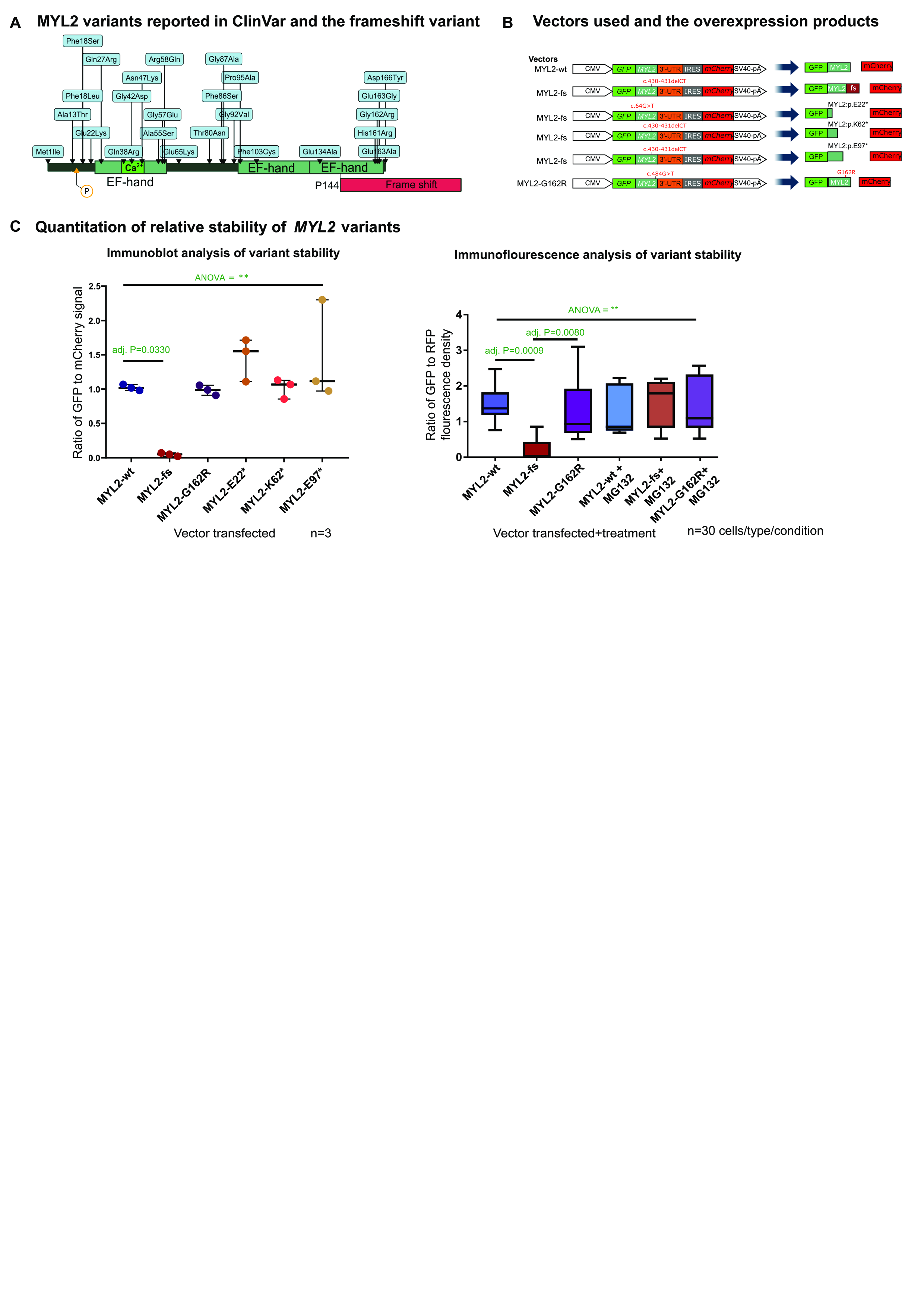

### Fig. S4

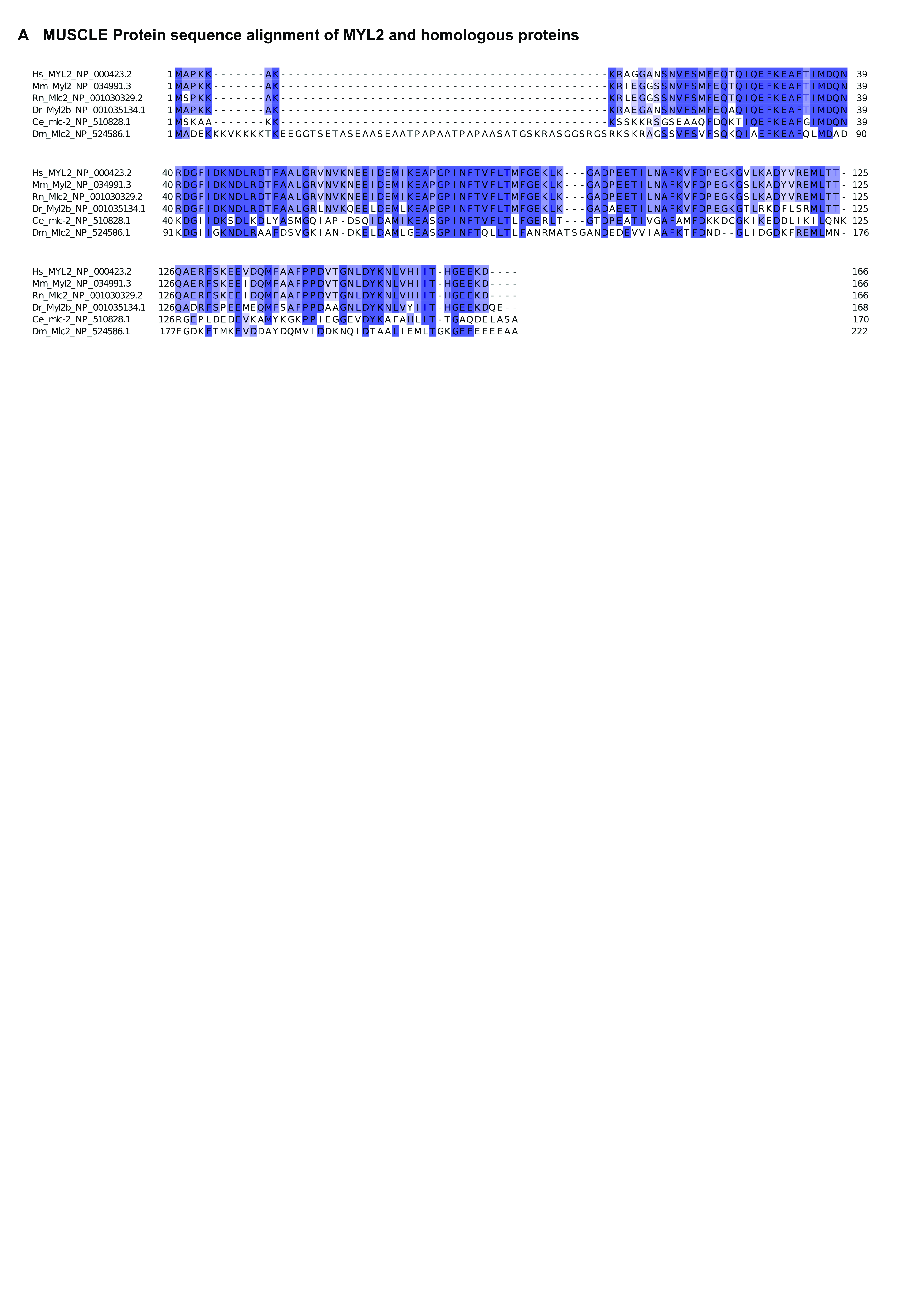
